# Cross-Platform Mechanical Characterization of Lung Tissue

**DOI:** 10.1101/271726

**Authors:** Samuel R. Polio, Aritra N. Kundu, Carey E. Dougan, Nathan P. Birch, D. Ezra Aurian-Blajeni, Jessica D. Schiffman, Alfred J. Crosby, Shelly R. Peyton

**Affiliations:** Department of Chemical Engineering, University of Massachusetts, Amherst; Polymer Science and Engineering Department, University of Massachusetts, Amherst

## Abstract

Published data on the mechanical strength and elasticity of lung tissue is widely variable, primarily due to differences in how testing was conducted across individual studies. This makes it extremely difficult to find a benchmark modulus of lung tissue when designing synthetic extracellular matrices (ECMs). To address this issue, we tested tissues from various areas of the lung using multiple characterization techniques, including micro-indentation, small amplitude oscillatory shear (SAOS), uniaxial tension, and cavitation rheology. We report the sample preparation required and data obtainable across these unique but complimentary methods to quantify the modulus of lung tissue. We highlight cavitation rheology as a new method, which can measure the modulus of intact tissue with precise spatial control, and reports a modulus on the length scale of typical tissue heterogeneities. Shear rheology, uniaxial, and indentation testing require heavy sample manipulation and destruction; however, cavitation rheology can be performed *in situ* across nearly all areas of the lung with minimal preparation. The Young’s modulus of bulk lung tissue using microindentation (1.9±0.5 kPa), SAOS (3.2±0.6 kPa), uniaxial testing (3.4±0.4 kPa), and cavitation rheology (6.1±1.6 kPa) were within the same order of magnitude, with higher values consistently reported from cavitation, likely due to our ability to keep the tissue intact. Although cavitation rheology does not capture the non-linear strains revealed by uniaxial testing and SAOS, it provides an opportunity to measure mechanical characteristics of lung tissue on a microscale level on intact tissues. Overall, our study demonstrates that each technique has independent benefits, and each technique revealed unique mechanical features of lung tissue that can contribute to a deeper understanding of lung tissue mechanics.

## INTRODUCTION

Lung tissue is highly elastic and mechanically robust over hundreds of millions of respiratory cycles. In order to properly ventilate the alveoli to facilitate gas exchange, it must maintain a delicate balance between strength and compliance to allow for these repeated, massive expansions. Lung parenchyma is the area of the lung that is involved with gas exchange, including the alveoli and smaller bronchioles, but excludes the large, cartilaginous bronchi. Lung parenchyma derives its mechanical integrity from the ECM, which is primarily composed of elastin, laminin, and collagen (Maksym and Bates 1997, Ito, Ingenito et al. 2005, Suki, Ito et al. 2005, Nichols, Niles et al. 2013). Others have shown that these structural proteins contribute to the mechanical properties of tissues (Liu and Tschumperlin 2011, Suki, Stamenovic et al. 2011).

Many research groups have measured and modeled the mechanical properties of the lungs with *in situ* and *in silico* techniques (Stamenovic 1990, Suki 2014), such as atomic force microscopy (AFM), uniaxial testing, and rheology; however, no group has presented data that directly compares these methods. Reports of lung parenchyma Young’s moduli range locally from ~2kPa (Liu, Mih et al. 2010) to greater than 15 kPa (Melo, Garreta et al. 2013) depending on the health of the tissue, the technique applied, and location of the measurement. This is an important point, as changes in stiffness by just a few kPa affects the phenotype of lung fibroblasts (Asano, Ito et al. 2017). The fact that the reports of lung moduli varies larger than changes sensed by cells makes it challenging for researchers in the biomaterials community to derive mechanobiological relationships, limiting applications in regenerative medicine and pathophysiology.

To determine the modulus of intact lung parenchyma, most procedures have traditionally applied pressure to excised lungs (Lai-Fook and Hyatt 2000, Dai, Peng et al. 2015), or used MRI (Mariappan, Kolipaka et al. 2012) or spirometry to determine their properties non-invasively (Gibson and Pride 1976). MRI yields shear stiffnesses ranging from 0.81 to 3.2kPa depending on the degree of inflation (Goss, McGee et al. 2006). Ultrasound is another non-invasive technique that has reported moduli from 752 to 954 Pa (Zhang, Qiang et al. 2011). Indentation, uniaxial extension, and other direct, local measurement devices are precise, but typically result in damage to the tissue structure from routine sample preparation. In contrast, the drawbacks to non-invasive methods are that they are very expensive and do not currently have the resolution to capture mechanical information on the sub-centimeter scale. These local heterogeneities could be an important factor for bioengineered tissue mimics given the lung’s heterogeneous composition.

Even still, many of the techniques measuring local mechanical properties are conducted on excised tissue samples. It is very difficult to determine local mechanical properties on small regions of the lungs without removing them, unless one uses a complex and expensive techniques, such as ultrasound (Zhang, Osborn et al. 2016) or MRI elastography (Mariappan, Kolipaka et al. 2012). Therefore, a simple, direct technique that would not require destructive preparation would be optimal. One such technique that has yet to be applied to lung tissue is cavitation rheology (Zimberlin, Sanabria-DeLong et al. 2007, Jansen, Birch et al. 2015), which does not require destructive sample preparation, is relatively inexpensive, and is portable. By not requiring significant handing or manipulation of the samples, one can perform *in situ* measurements of the lung tissue in as close to its relaxed state as possible without tissue destruction.

In this study, we describe the application of cavitation rheology to lung tissue, and we attempt to unify tissue modulus measurements by comparing cavitation rheology to other traditional tissue mechanics approaches (Zimberlin, Sanabria-DeLong et al. 2007). We compared cavitation to popular techniques in the field: micro-indentation, uniaxial testing via Instron, and SAOS with a parallel plate rheometer (Figure 1). By performing these techniques on the same samples, we provide a direct comparison across measurement modalities. This data provides important information that the field can use to recapitulate the lung modulus with tissue models, and provides insight into local mechanical properties of the intact lung.

**Figure 1.**
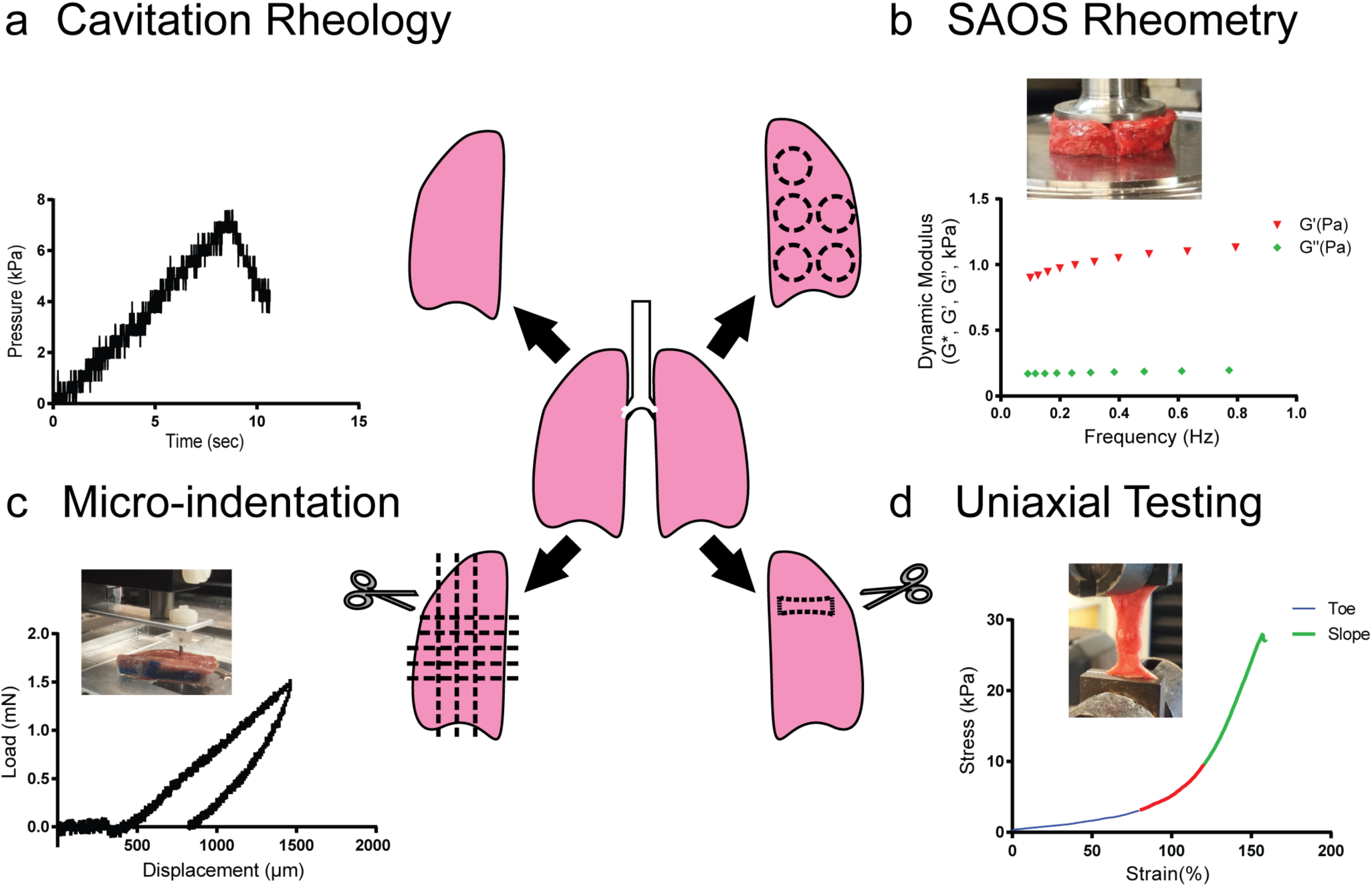
Methods Used to Quantify Lung Modulus. Lungs from **12** animals were tested using multiple methods. For each method, an example force-displacement or force-time curve is shown for a typical tissue sample, as well as an inset image of the tissue sample being actively tested. (a) Cavitation rheology uses a small needle inserted into an intact tissue, wherein a sensor measures the pressure required to cavitate a bubble within a tissue. This cavitation pressure was then input into a model to find a tissue compliance and an effective modulus. (b) SAOS requires defined, excised samples and applies shear across a range of oscillatory frequencies, resulting in a dynamic modulus, storage modulus (G’), and loss modulus (G”), which can be related to the Young’s modulus via the Poisson’s ratio (*v*). (c) Micro-indentation requires excised samples with defined thicknesses and applies small compressive forces, results in a force-displacement curve, the slope of which resulted in a Young’s modulus. (d) Uniaxial testing applies tension to a prepared, excised sample, resulting in a stress-strain curve, the slope of which relays a Young’s modulus. Due to the heterogeneity of the ECM in structure and composition, lung tissue has two different regimes: an initial toe region, and a steeper, sloped region.

## MATERIALS AND METHODS

### Lung Tissue and Sample Preparation

Porcine lungs were acquired from Research 87 (Boston, MA) and immediately placed on ice for transport. The tissue was mechanically tested as soon as possible (within 3–4hr) post-slaughter. After removal from ice, the lungs were thoroughly rinsed with 1x phosphate buffered solution (PBS; Sigma Aldrich, St. Louis, MO), divided into 1 × 1 in^2^ sections to facilitate localization and uniformity of measurements across techniques and between lung samples (Supplementary Figure 1). In the case of cavitation rheology, the lungs remained in the plastic wrap for the duration of the testing to prevent dehydration. For uniaxial testing, SAOS, and micro-indentation, the grids were used as guides to divide samples within the lung, and dehydration was minimized by placing samples into 6-well plates with water in the regions between the wells.

In instances where we compared fresh to frozen tissues, sample sections were flash frozen in liquid nitrogen (LN2) and stored at −80°C, or placed in a 6-well plate and frozen at −80°C, or placed in optimal cutting temperature (OCT) medium (Fisher Scientific, Hampton, NH) in a weighing dish, which was exposed to LN2 until the sample was completely frozen after initial testing. Samples frozen in OCT medium were defrosted prior to measurement and rocked on a shaker with 1x PBS at 22°C for 20 min to completely thaw.

### Baseline PEGDMA/HEMA Hydrogel Preparation

For the synthetic testing material, we used a poly(ethylene glycol) (PEG) hydrogel biomaterial consisting of a 1:10 molar ratio of PEG dimethacrylate (PEGDMA, Sigma-Aldrich, St. Louis, MO) to 2- hydroxyethylene methacrylate (HEMA, Sigma-Aldrich). For a given vol% PEGDMA in a 1:4 200 proof ethanol (Pharmco-Aaper, Brookfield, CT): dimethylsulfoxide (DMSO, VWR, Radnor, PA) solution, we added HEMA and 4 vol% of the UV-sensitive free radical initiator Irgacure 2959 (BASF, Florham Park, NJ). The gels were UV treated for at least 30 min and swollen overnight in 1:4 ethanol: DMSO solution.

### Cavitation Rheology

Cavitation was carried out on porcine lungs using a custom-built instrument as described previously (Zimberlin, Sanabria-DeLong et al. 2007, Jansen, Birch et al. 2015) (Figure 1a). Briefly, the instrument is composed of a syringe pump (Nexus 6000; Chemyx Inc., Stafford, CT), pressure sensor (PX26-005DV; Omega, Stamford, CT), and a syringe needle connected to a DAQ card to record the pressure over the course of the experiment. Needles with varying gauges of 16, 18, 22, and 26 (1.194, 0.838, 0.413, 0.260 mm inner diameters, respectively) (Hamilton Company, Reno, NV) were inserted into the lung until they pierced the viscera pleura. The needles had beveled tips to decrease the stress on the needle and to pierce the lung. Cavitation rheology on PEG hydrogels was performed using flat needles with varying gauges of 24, 27 and 30 (0.311, 0.210, 0.159 mm inner diameters, respectively) (Hamilton Company, Reno, NV).

Air was injected at a rate of 5000 μL/min through the needle via the syringe pump and the injection continued until there was a significant pressure drop, signifying a cavitation event. The strain rate was calculated based on the air injection rate, and the modulus was calculated using the following equation (Hutchens and Crosby 2014, Hutchens and Crosby 2014):

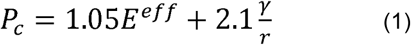

Where *P_c_* is the cavitation pressure, *E^eff^* is the effective Young’s modulus, *γ* is the effective surface tension between tissue and the injected air, and *r* is the needle radius. The coefficients, as well as the general form of this equation, were determined for an ideal material described by a neo-Hookean constitutive relationship; and, we have previously applied it to biological tissues (Cui, Hee Lee et al. 2011, Chin, Freniere et al. 2013, Jansen, Birch et al. 2015). The y-intercept of the cavitation pressures vs. 1/*r* was used to determine the effective Young’s modulus following equation (1). A minimum of 3 needles was used to calculate the cavitation pressure of a particular 1 × 1 cm^2^ region of the lung (as depicted in Supplementary Figure 1). Data analysis was carried out using MATLAB (Natick, MA) after filtering with a Butterworth filter.

### Small Amplitude Shear Oscillation

SAOS (Figure 1b) was performed using an oscillatory test on a Kinexus Pro Rheometer (Malvern Instruments, UK) with a 20 mm diameter flat plate geometry and a gap between 1.5–2.0 mm. Samples were obtained from portions of the lung closer to the periphery and in plane with the cutting surface (Supplementary Figure 1) to avoid large bronchioles or cavities within the lung tissue. Excess material was trimmed with a scalpel and surgical scissors. A solvent trap was used to prevent the samples from dehydrating over the duration of the experiment and the plate was maintained at either 25°C or 37°C to test the tissue sensitivity to temperature. A strain sweep concluded that a strain of 0.5% would place the measurements within the linear regime of the tissue (Figure 4a). Oscillatory frequency sweeps were conducted between 0.1 and 1.0 Hz based on the shear rate and the total deformation of the material. Since the tissue was deflated for all measurements, the alveolar pressure was assumed to be zero. The Young’s modulus (E) was calculated at a frequency of 0.1 Hz using the complex shear modulus (G*) and a Poisson’s ratio (v) of 0.42 according to the equation (Butler, Nakamura et al. 1986):

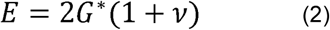

### Micro-Indentation

Indentation testing was performed using a custom indenter as described by Chan et al. (Figure 1c) (Chan, Smith et al. 2008). A flat cylindrical probe with a radius of 0.5 mm was made from tool hardened steel (McMaster-Carr, Elmhurst, IL) and attached to a flat, deformable, 0.40 mm wide cantilever. The indentation rate was kept at a constant 20 μm/s for each sample and indented to a maximum force of 2 nN as measured by a Honeywell Sensotec (Columbus, OH) capacitive force transducer. The strain rate was calculated based on the rate of indentation, normalized by the thickness of the sample. The transducer was in series with a nanoposition manipulator (Burleigh Instruments, Rochester, NY, Inchworm Model IW-820) that controlled displacement and the rate of displacement. The force (*F*) and displacement (*δ*) measurements were recorded with a custom LabVIEW program (National Instruments, Austin, TX). The material compliance (*C*) was measured over the linear region after coming into contact with the surface using the following equation:

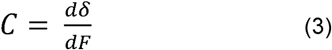

The deflection of the cantilever was taken into consideration by subtracting the deflection required to produce the given force from the displacement indicated. Up to 3 measurements were conducted per sample on areas that were not cartilaginous or containing bronchioles nor close to the edge of the sample. The Young’s modulus (*E*) was calculated using equation 4 after taking into account the height of the sample (*h*), the Poisson’s ratio of 0.42 (*v*) and the radius of the circular indenter (*a*), assuming the material is elastic, according to Shull *et al*. (Shull, Ahn et al. 1998).

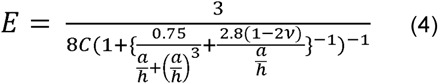

### Uniaxial Testing

Uniaxial testing was carried out using a model 5564 Instron (Instron, Norwood, MA) with a 50 N load cell in a manner similar to the procedures found in O’Neill et al. (Figure 1d) (O’Neill, Anfang et al. 2013). It is relevant that we are measuring engineering stress and assuming that the sample is nearly incompressible and isotropic over small strains. Samples were cut using surgical scissors into approximately 2.7 cm × 1 cm × 2 mm sections. Samples were taken from sections with minimal cartilaginous regions (Supplementary Figure 1) and strained at a rate of 1% of the starting length per second until failure. The strain rate was calculated by normalizing the global displacement rate by the total stretch length.

### Statistics

Statistics were performed using Prism V6.05 and compared using two-tailed Student’s t-tests where appropriate. Error bars show standard error unless noted. An asterisk denotes p < 0.05.

## RESULTS

### Freezing Lung Tissue Causes Small but Detectable Reduction in Young’s Modulus

In order to conduct efficient studies on the lung tissue, we needed to determine if freezing the samples to measure at a later date would damage them. Freezing the lung samples after slaughter is the most common method for labs to preserve decellularized (Nonaka, Campillo et al. 2014) and native tissues (Graham, Hodson et al. 2010). Freezing or embedding into agarose (Booth, Hadley et al. 2012) is often the only way AFM sections can be prepared for indentation when thin samples are required. It has been suggested that freezing the lung tissue using LN2, at −80°C, or using OCT medium to freeze samples would not significantly impact these mechanical properties (Figure 2). However, our results demonstrate a small, but detectable, trend in decreasing modulus after freezing as measured by micro-indentation, across all the freezing methods attempted (Figure 2).

**Figure 2.**
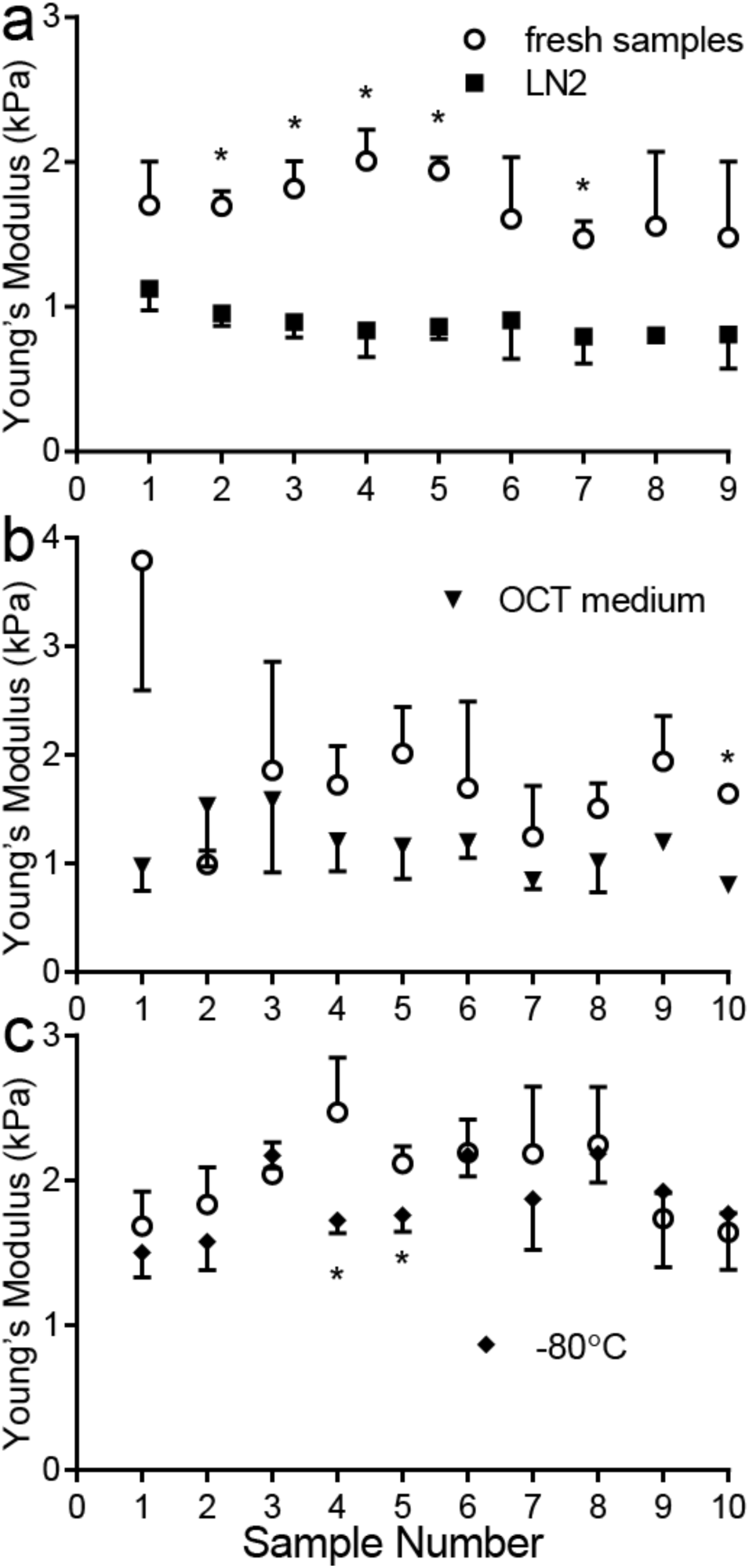
Mechanical Properties of Fresh vs. Frozen Tissue using Micro-Indentation. Fresh lung samples were tested using micro-indentation and then frozen using either (a) Liquid Nitrogen (LN2), (b) OCT medium with LN2, or (c) slowly frozen to −80°C, and subsequently re-tested. Samples marked with * are statistically significantly different after freezing, compared to the original fresh tissue specimen, (p<0.05) as found using a Student’s t-test. Error bars are shown in only one directly for clear visualization of the data.

When comparing individual samples, the modulus of the tissue was typically lower when subjected to freezing compared to freshly isolated tissue, and this comparison was statistically significant most often in the LN2 freezing process. However, when data was compiled across the entire population of tissues for each technique, comparisons were significant in cases where tissues were frozen in OCT medium or LN2 (p<0.05), but not when frozen at −80°C (p>0.05). The average Young’s modulus as measured by microindentation of the fresh lung samples was in general lower, regardless of the freezing technique. In OCT medium, the Young’s modulus of the samples dropped from 1.8±0.8 kPa to 1.2±0.3 kPa (N=10). Similarly, directly freezing the samples in LN2 caused the stiffness to drop from 1.7±0.2 kPa to 0.9±0.1 kPa (N=9). Freezing samples at −80°C caused a stiffness decrease from 2.0±0.3 kPa to 1.9±0.1 kPa (N=10), but this was not statistically significant. The subsequent tests in this paper were all carried out directly after slaughter without freezing to avoid any effects that freezing could have on the outcome of the testing. Although preservation of individual samples could be done, there was particular concern about how thawing an entire lung would affect sample integrity, particularly for cavitation rheology.

### Lung Tissues can be Characterized with Multiple Mechanical Techniques, in any Order

Given possible constraints in equipment availability when tissues arrive at a lab, we sought to determine if individual samples could be tested with multiple techniques, and if certain techniques were sufficiently destructive to prohibit subsequent, accurate testing by other methods. Our results show that when SAOS was performed before indentation, the aggregate Young’s Modulus as measured with microindentation slightly increased to 2.2±0.1 kPa from 2.0±0.1 kPa, but this difference was not statistically significant (Figure 3). The procedures used to measure micro-indentation did not change the modulus as pre-micro-indentation measured shear modulus was 1.03 ± 0.04 kPa and post-micro-indentation was 1.09±0.04 kPa.

**Figure 3.**
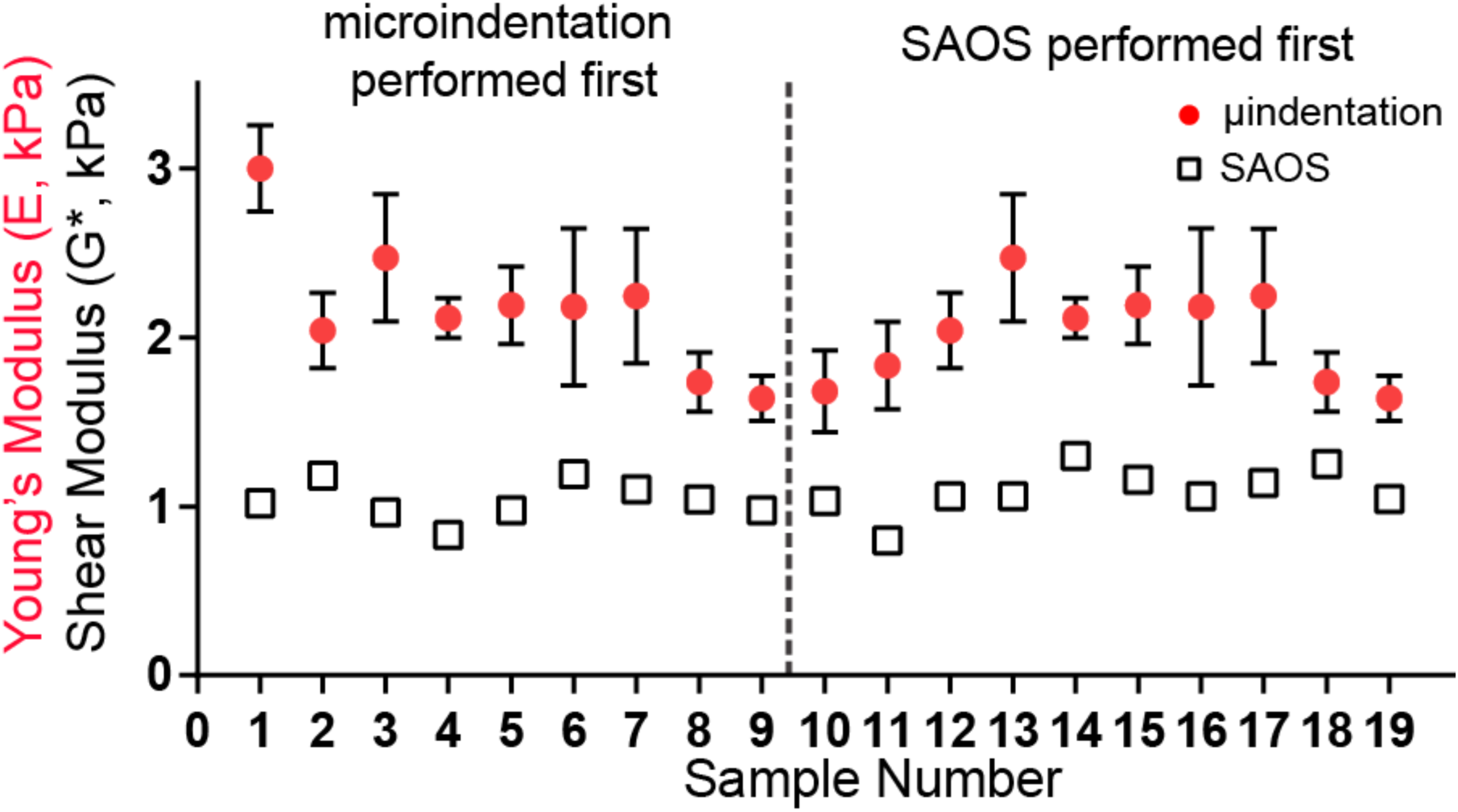
SAOS and Micro-Indentation Moduli Do Not Depend on Testing Order. The Young’s modulus measured with micro-indentation before (left of dashed line) or after (right) SAOS did not affect the measured sample modulus. SAOS samples could be tested once (N=1), while micro-indentation samples were indented multiple times (N=3) to obtain the mean and standard deviation.

### Temperature Variations do not Significantly Affect SAOS of Lung Samples

For SAOS, the samples were tested using a frequency sweep at 25° and 37°C to determine the linear regime of the samples (Figure 4a). Our results found a linear strain response at less than 1% strain and further testing was confined to the linear viscoelastic region. At these low strains, we conducted a frequency sweep to verify that there was minimal effect on the modulus as we varied the frequency. Furthermore, we found that increasing the temperature from 25°C to 37°C resulted in no measurable differences between the samples’ moduli (Figure 4b).

**Figure 4.**
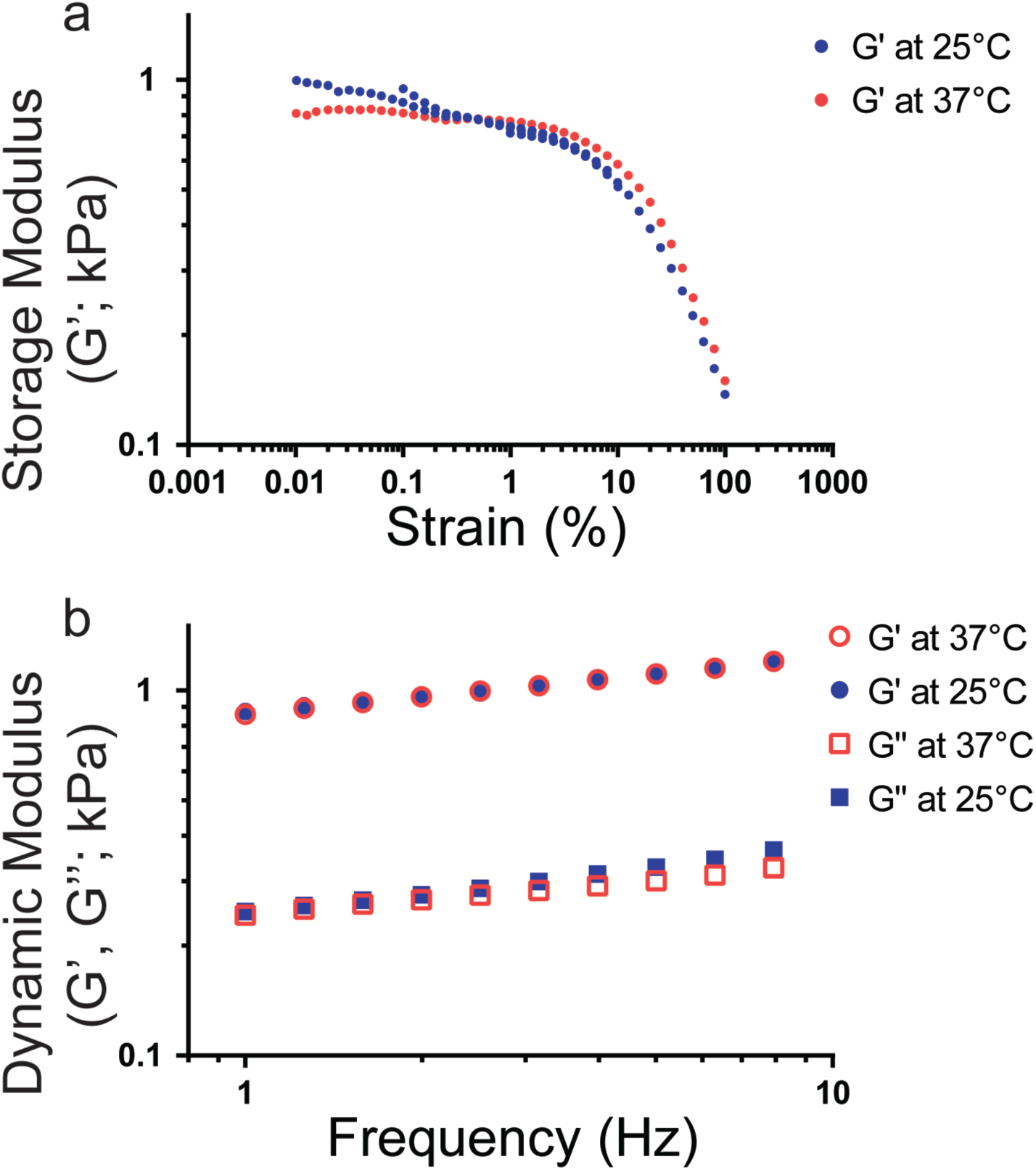
Lung Tissue Modulus Temperature Dependence as Measured by SAOS. (a) A lung sample was subjected to a strain sweep from **0.01%** to **100%** at both 25°C and 37°C. Multiple runs were conducted on the same sample to construct the plot. (b) A sample of lung tissue was tested at 25°C and 37°C to determine the effect on temperature variation. No statistically significant differences were noted in the storage modulus (G’) or loss modulus (G”) as a function of temperature.

### Comparative Measurements Show an Increased Modulus Using Cavitation Rheology

We directly compared measurements made using micro-indentation, SAOS, and cavitation rheology across several lung tissue samples, as well as a PEG hydrogel system (Figure 5). Cavitation rheology reported the stiffest moduli, with a mean of 6.1±1.6 kPa, while micro-indentation testing reported the most compliant moduli of 1.4±0.4 kPa. SAOS reported a modulus of 3.3±0.5 kPa. SAOS and micro-indentation measurements were generally similar to the order of magnitude of the “toe region” from uniaxial tension (3.4±0.4 kPa), where there is a shallow slope, associated with a low Young’s modulus (Figure 1d).

**Figure 5.**
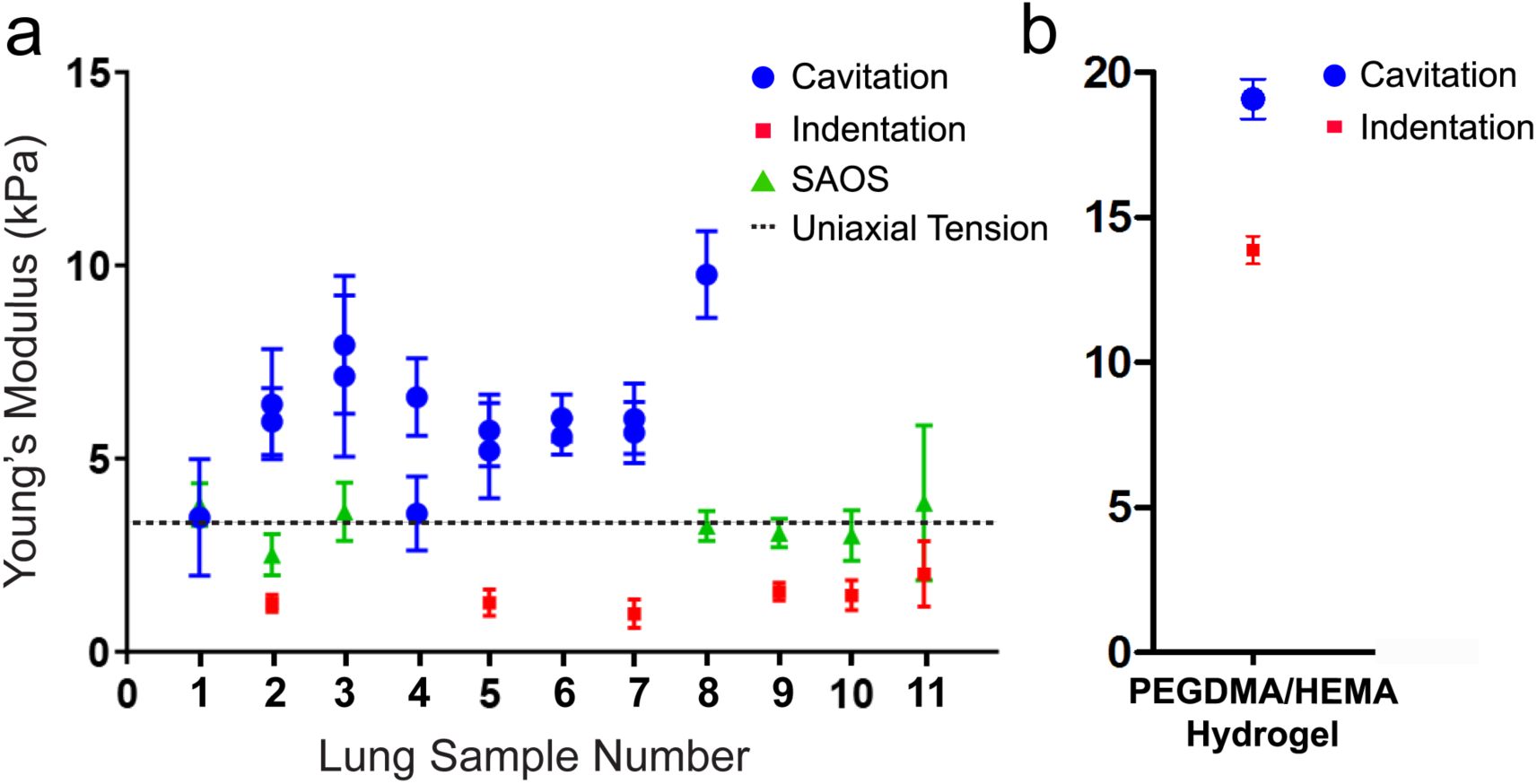
Comparison of Young’s Modulus of Lung Tissue and Baseline Hydrogel across Different Mechanical Testing Techniques. (a) The Young’s modulus of the parenchymal region of the lungs were determined by each method described: cavitation rheology (blue), micro-indentation (red), SAOS (green) and uniaxial tension (dashed line). Each data point represents averaged data across samples using the method indicated for a single lung lobe. Lobes within sets of lungs were paired to generate up to four data points per lung set (sample number). (b) The Young’s modulus of a 4 vol% PEGDMA/HEMA hydrogel was determined by cavitation rheology (blue), and micro-indentation (red). Each data point represents averaged data across several different hydrogel samples (N=14 for cavitation, N=11 for indentation).

Cavitation rheology showed significant variation in moduli across samples. It was only when averaging each needle diameter for the whole lung (Supplementary Figure 2) that we were able to obtain enough data to represent the Young’s modulus. Multiple sample measurements in the same gridded area reflected the tissue heterogeneity.

However, after correcting for differences in strain rates, we did observe improved correlations between SAOS and micro-indentation (Supplementary Figure 3). This correction was done by scaling the Young’s modulus using the power law fit:

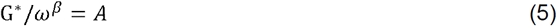
where *ω* is the relative frequency at which the test was conducted, *β* is a dimensionless parameter derived from SAOS (*β* = 0.11), and *A* is a proportionality constant relating the shear modulus and frequency. We found the proportionality constants for each test by fitting the SAOS samples to this power law model.

We then performed cavitation rheology on a baseline 4 vol% PEGDMA/HEMA hydrogel to determine whether variability in cavitation rheology testing was due to the method itself, or heterogeneities within and across lung tissue samples. In a synthetic PEG hydrogel, the measured critical pressures were extremely consistent, resulting in similarly consistent moduli (Supplementary Figure 3). Similar to the lung tissue results, cavitation rheology on the PEG hydrogels reported stiffer moduli (19.1±0.7 kPa) than microindentation (13.9±0.5 kPa, Figure 5).

Across the different characterization techniques applied here, the strain rates differed by up to an order of magnitude. These strain rate variations could have significantly affected the reported moduli across the different techniques. To determine the strain rate variance possible on a model hydrogel material, we measured the response of the PEGDMA/HEMA hydrogels to microindentation at four different strain rates (0.01 Hz, 0.03 Hz, 0.05 Hz, and 0.10 Hz). These strain rates encompassed the range applied by each individual technique (SAOS, indentation, cavitation, and uniaxial tension). No significant variation was seen in the moduli at the different strain rates (Supplemental Figure 4).

## DISCUSSION

The mechanical properties of the lungs, such as compliance, are integral to this organ’s biological functions. However, the lung is a structurally and mechanically heterogeneous tissue, with regions of high and low stiffness and porosity, which are difficult to comprehensively capture. Significant research has been performed to understand lung mechanics to fully grasp this complexity and in an attempt to correlate lung structure and compliance with organ-, tissue-, and cell-scale behaviors. One challenge in achieving this goal is that studies have used a variety of different techniques quantified the compliance of lung tissue: from indentation on small, excised samples to elastography on of whole lungs *in situ*. Therefore, one of our goals was to take the first step in unifying a variety of different methods, we directly compared various testing modalities to determine how the reported modulus of lung tissue varies across them given the inherent constraints for each method: differences in sample preparation, temperature, and region of tissue tested. This effort will enable the field to compare measured and reported values, using different methods to quantify a tissue modulus, and will help to create a more accurate hydrogel model for simulating *in* vivo-like compliance of the lung. In doing this comparative study, our most significant findings are that the modulus of lung tissue does vary depending on the testing modality used (though each reported value is within the same order of magnitude), the order of testing does not significantly affect the reported modulus, freezing tissue samples has a modest, though detectable effect on the reported modulus, and cavitation rheology is the only method capable of reporting a localized *in situ* modulus.

We obtained and tested samples as soon as possible post-slaughter (within hours, and tissues were kept hydrated). *In vivo* lung tissue is in a pre-stressed state (Suki, Stamenovic et al. 2011), therefore, some variation between our measurements and those obtained *in vivo*, if that were possible, would be expected. Practically speaking, many labs will not be able to use freshly obtained lung tissue, as we did here, and so we also quantitatively compared the effect of several freezing techniques on the lung tissue mechanical properties. We found a detectable decrease in Young’s modulus when flash freezing using LN2, but not when slow freezing to −80°C. The benefit is that these techniques did not require samples to be flash frozen (Graham, Hodson et al. 2010, Luque, Melo et al. 2013) or embedded in agar (Liu, Mih et al. 2010, Liu and Tschumperlin 2011) prior to measurement, which would have introduced additional confounding mechanical changes to the samples.

There is limited knowledge on nonlinear, viscoelastic, heterogenous, and anisotropic materials. Therefore, we employed techniques that are commonly used in the tissue mechanics field to approximate the linear and non-linear responses of lung tissue. From our measurements, we determined that microindentation, cavitation rheology, uniaxial extension, and SAOS tests were in line with some of the previously reported values for the lung (~5 kPa) (Suki and Bates 2008, Booth, Hadley et al. 2012, Luque, Melo et al. 2013). The experimental modulus values for lung tissue for micro-indentation (1.9±0.5 kPa), SAOS (3.2±0.6 kPa), and uniaxial testing (3.4±0.4 kPa) are based on measuring a modulus for the bulk material, however cavitation (6.1±1.6 kPa) measures the modulus on a length scale approaching that of the microstructures of the lung. We suggest that the information provided with these different measurement modalities can spur further interest in refining *in silico* mechanical models of the lung microenvironment.

Our second goal was to demonstrate a new technique for measuring the modulus of lung tissue at similar length scales to that of microstructure heterogeneities within tissue. Cavitation rheology was the least destructive technique we applied in that we did not need to excise any tissue to perform the testing; however, the range of measurements on individual samples, as well as the average values across multiple lung tissues, was detectably higher in comparison to values obtain from the other approaches used here. The higher modulus of cavitation may be the result of ECM fiber reorientation during extension of the material. In support of this speculation, we observed strain-hardening with uniaxial extension, though not with SAOS or micro-indentation. The nonlinearity of the uniaxial testing can be largely attributed to the alignment and extension of collagen, which is a main contributor of mechanical strength to tissue (Suki, Stamenovic et al. 2011). It is possible that cavitation rheology is more sensitive to the mechanical response at larger strains for tissues with nonlinear properties, but more fundamental studies of cavitation dynamics are required to understand these responses. It is difficult to extract the contributions of the individual ECM proteins in the tested tissue using these techniques; however, it would be interesting to explore a similar gamut of modalities to find how the moduli compare. Modifications to the cavitation model could be extracted from direct observation of deformations under pressure mimicking *in* vivo-like conditions or through degrading various ECM components (Black, Allen et al. 2008). In previous work, our lab found that bone marrow modulus also agrees cross-platform, however without compensation for strain rate, cavitation rheology reports a higher modulus as compared to SAOS and micro-indentation (Jansen, Birch et al. 2015). Due to the small diameters of the needles, cavitation testing was more localized than the other techniques we used. Higher variability from cavitation compared to indentation and SAOS may have been due to fact that bronchiole-free areas were selected in the excised tissues for indentation and SAOS (Supplementary Figure 1). It is likely that in addition to sample selection, cavitation rheology is more sensitive to local heterogeneities in the tissue, much like AFM.

To confirm the trend that cavitation rheology testing results in a higher modulus value than other techniques, we tested a PEG hydrogel across the multiple mechanical methods. This PEG hydrogel can have a wide range in elastic moduli, which depends on the molecular weight of the polymer and the concentrations of the crosslinker (Peyton, Raub et al. 2006, Killion, Geever et al. 2011). After testing multiple conditions (data not shown), we chose gels with 4 vol% PEGDMA, as it had a stress-strain response and a Young’s modulus similar to the lung tissue samples (Figure 5). We used the PEG gels to determine if the consistently higher moduli reported by cavitation rheology found in the lung samples, and previously in bone marrow tissue (Jansen, Birch et al. 2015), was also found in a polymer hydrogel. When we compared cavitation rheology to indentation, we found that cavitation reported a higher modulus than indentation, consistent with the measurements in tissues (Figure 5b). Furthermore, we have previously published that cavitation rheology reports a higher modulus compared to other testing methods in synthetic hydrogel systems (Zimberlin, Sanabria-DeLong et al. 2007, Zimberlin and Crosby 2010). Because cavitation rheology is measuring the modulus of a material on length scales in the tens of microns, it is possibly sensitive to the mechanical contribution of local lung tissue microstructures. This is one possible explanation for more consistent results in the polymer hydrogels compared to the lung tissue samples (Supplemental Figure 4). Indentation, SAOS, and uniaxial strain were potentially not sensitive enough to sense the mechanical contribution of microstructures, or handling and excision resulted in significant tissue damage that caused a decrease in the observed moduli.

There are other factors to consider as contributors to variation in modulus measurements, such as the strain rate at which the test is carried out, the contributions of stiff elements of the lungs that cannot be seen during testing, such as bronchioles near the testing area, variation in alveolar size, and potential contributions due to porosity (Litzlbauer, Korbel et al. 2010, Jansen, Birch et al. 2015). We applied a strain rate of 0.01Hz in uniaxial extension, which was calculated by normalizing the global displacement rate by the gauge length. Cavitation rheology testing was performed at a strain rate of 0.028Hz, estimated based on the air injection rate. Indentation tests had a compressive strain rate of 0.04Hz, based on the rate of compression, normalized by the sample thickness. The strain rates applied in SAOS varies from 1 to 10 Hz based on the shear rate and the total deformation of the material. To determine how strain rate contributed to differences in the reported moduli, we varied the strain rate of indentation on PEG hydrogels between 0.1 and 1 Hz, encompassing the full range of strain rates covered by the different testing methods. We observed that the modulus was independent of the strain rate, at least across this range, suggesting that differences in lung moduli reported by each of these individual measurements was not due to strain rate differences.

Beyond strain rate differences between measurement methods, another possible contribution to the differences between measurements is the model used to interpret cavitation rheology. The model for relating cavitation pressure to the Young’s modulus used in this study was developed for flat needles, and our use of beveled needles to reduce tissue damage at the surface could have impacted the measurements, as there is a difference in the needle diameter at the opening versus the actual diameter of the needle’s shaft in the beveled tip geometry (Equation 1). In addition, due to the opacity of the tissue, we could not visually confirm if fracture or cavitation occurred within the tissue (Kundu and Crosby 2009, Zhang, Osborn et al. 2016). Since the model applied to the tissue is intended for non-porous, elastic materials, adjustments to this model could allow for a more detailed understanding of the relationship between cavitation pressure and the lung’s mechanical properties.

With these differences acknowledged, we still confirm that cavitation rheology provides a consistent measurement of mechanical properties on a tens of microns scale, providing a method for characterizing the localized heterogeneities within lung tissue. This attribute is significant since this length scale is likely more pertinent to how cells interact with their local environment. Therefore, measuring the modulus of tissues and biomaterials with cavitation provides significant advantages not only in characterizing mechanical properties of lung but also when designing materials to represent lung tissue.

## ACKNOWLEDGMENTS

We would like to acknowledge Mike Imburgia, Shruti Rattan, and Lauren Jansen for their help with uniaxial testing, micro-indentation, and cavitation rheology. Igor Kheyfets helped with preparing the protocols for cavitation testing. S. Polio, S. Peyton, and D. Aurian-Blajeni were funded by an NIH New Innovator Award (1DP2CA186573-01) and a grant from the NSF (DMR-1454806). C. Dougan, A. Crosby, and S. Peyton were supported by a BRC award from the ONR (N00014-17-1-2056). S. Peyton is a Pew Biomedical Scholar supported by the Pew Charitable Trusts, and was supported by a Barry and Afsaneh Siadat Faculty Development Award. N. Birch and J. Schiffman were partially supported by the Professor James M. Douglas Career Development Faculty Award and the Armstrong Fund for Science. A. Crosby was funded by a grant from the NSF (DMR-1304724).

The authors declare no conflicts of interest for this work.

---

**Supplementary Figure 1.**
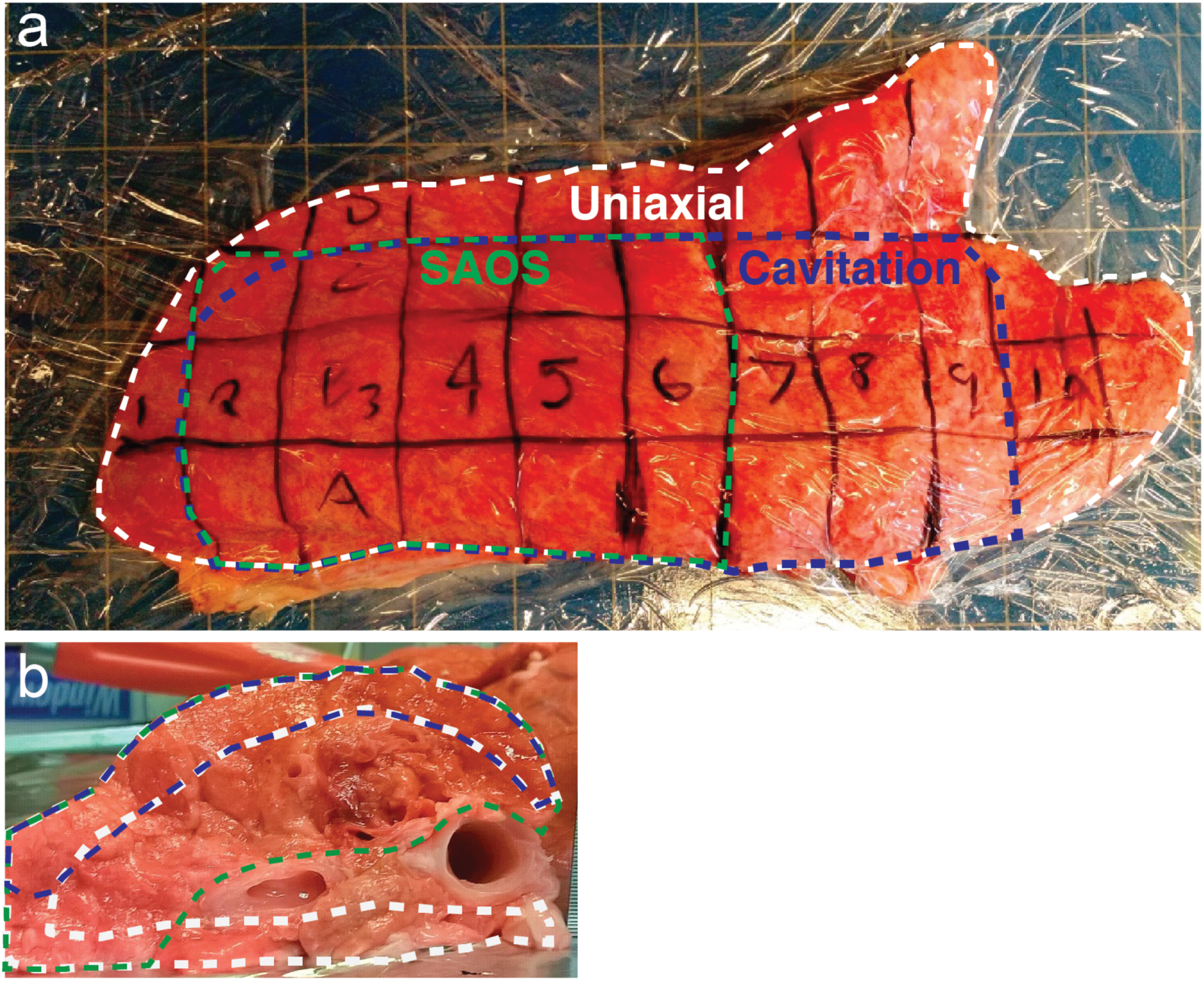
Lung Gridding and Tested Tissue Areas for each Technique. (a) Shown is an example of the grid system used for marking the lung tissues prior to testing and/or excision. The lungs were wrapped in plastic wrap and a grid consisting of 1 in × 1 in squares was drawn onto the wrap in order to help identify lung samples. (b) This image shows the inner structure of the lung (near column 4 in (a)), which is extremely heterogeneous and cartilage-rich that had to be avoided during tissue excision for SAOS, microindentation, and uniaxial tension. The colored, dashed lines denote the tissue areas used for each technique. Micro-indentation can be taken from anywhere within the lung, but there are restrictions on where samples can be taken for other tests such as cavitation (blue), uniaxial testing (white) and SAOS (green). Samples were excised as outlined in the dashed lines in (a) and (b) for each method, avoiding the cartilaginous bronchioles.

**Supplementary Figure 2.**
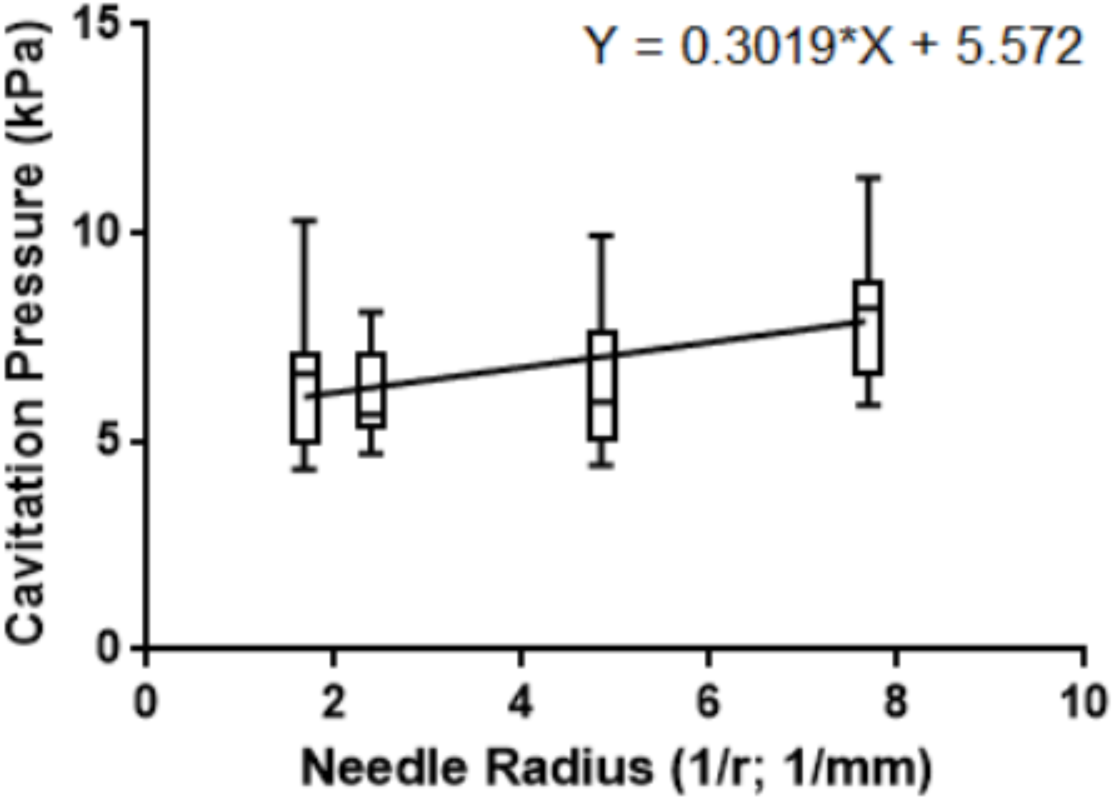
Representative Cavitation of a Porcine Lung. The Young’s modulus of the sample can be found by fitting a line to the data and finding the intercept (5.6±0.5 kPa). Error bars represent the standard deviation. The intercept can be used to determine the elasticity of the material.

**Supplementary Figure 3.**
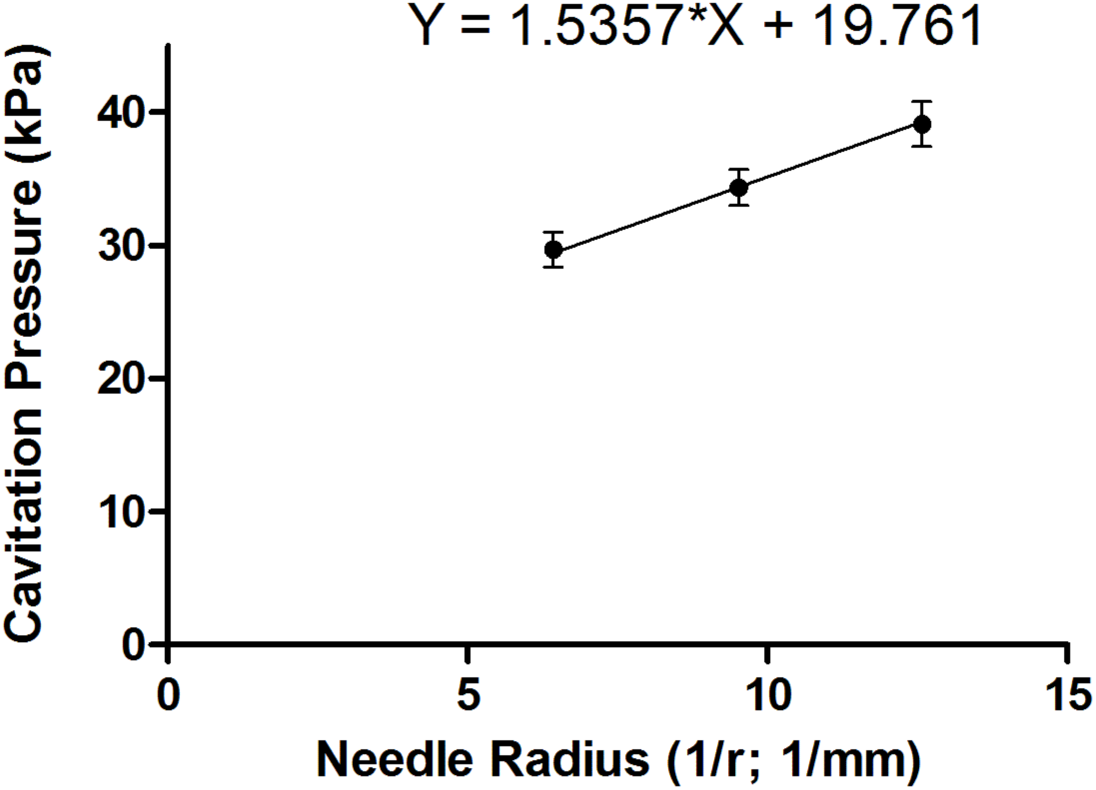
Representative Cavitation of a 4 vol% PEGDMA/HEMA Hydrogel. The Young’s modulus of the sample was determined by fitting a line to the data for cavitation pressure versus needle radius (N=6), and finding the y-intercept (19.8±0.6 kPa). Error bars represent the standard deviation. The intercept is the effective modulus of the material.

**Supplementary Figure 4.**
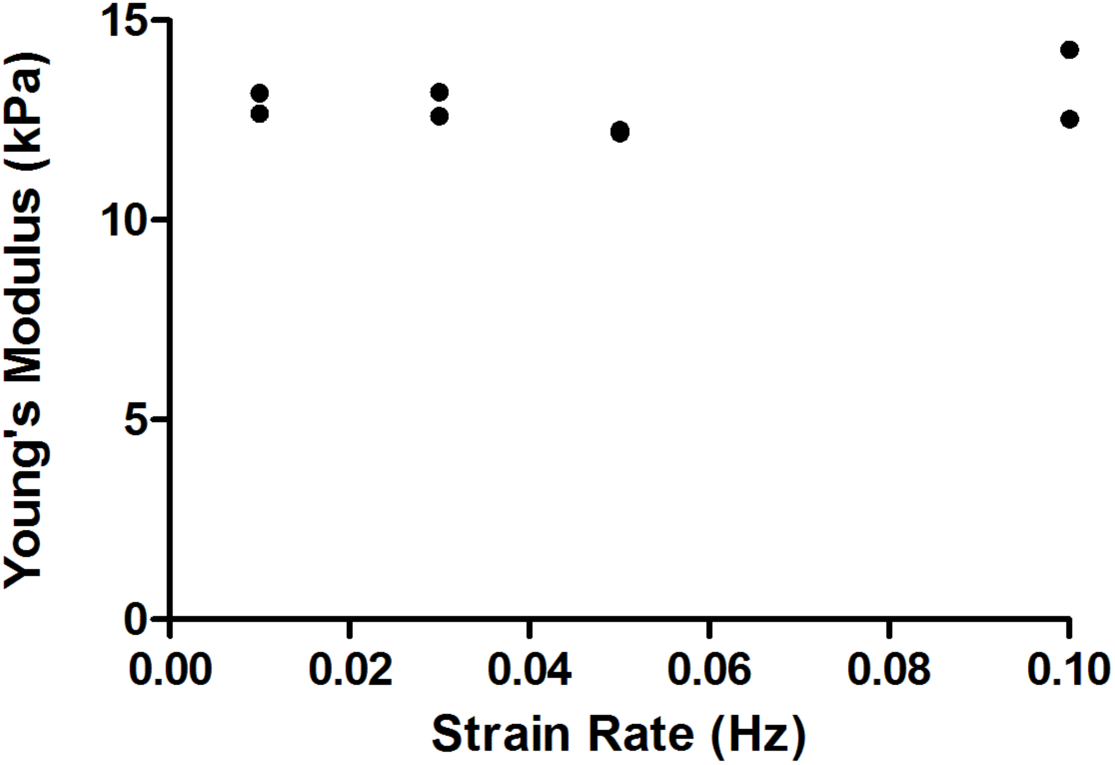
Strain Rate Dependence of Hydrogel Biomaterial Modulus. The Young’s moduli of the baseline material 4 vol% PEGDMA/HEMA hydrogels swollen in ethanol/DMSO solution, were determined by micro-indentation as a function of strain rate. The strain rate was varied from 0.01 Hz to 0.10 Hz: 0.01 Hz, 0.03 Hz, 0.05 Hz and 0.10 Hz. Micro-indentation was performed across 2 samples for each strain rate, and the Young’s moduli were not statistically significantly different from each other, as determined by a Student’s t-test.

**Supplementary Figure 5.**
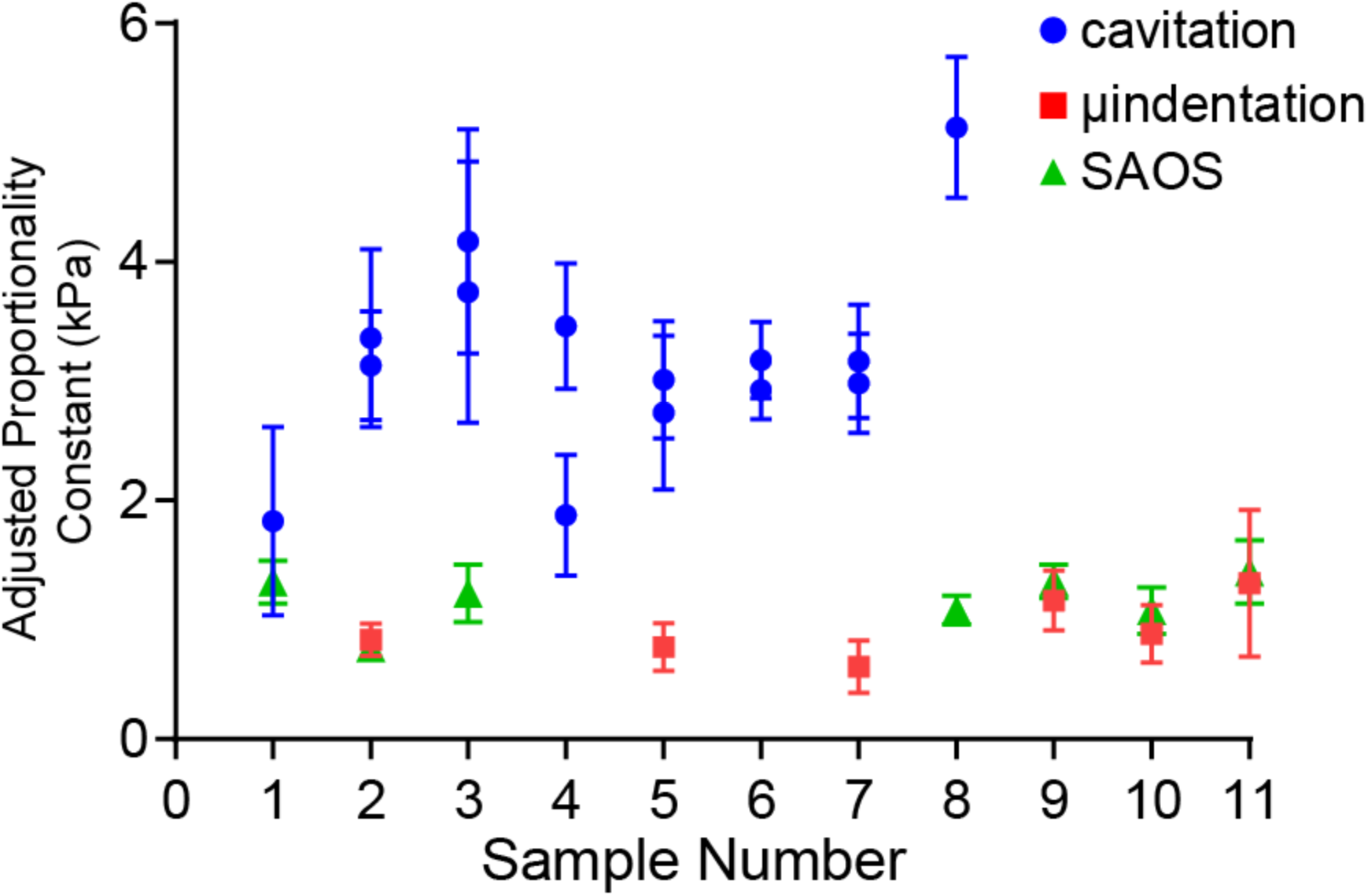
Modulus Proportionality Constant for the Techniques. The Young’s moduli were adjusted based on the power law to compensate for the differences in the frequency of the tests.

